# A large-scale population study of early life factors influencing left-handedness

**DOI:** 10.1101/305425

**Authors:** Carolien G.F. de Kovel, Amaia Carrión-Castillo, Clyde Francks

## Abstract

Hand preference is a conspicuous variation in human behaviour, with a worldwide proportion of around 90% of people preferring to use the right hand for many tasks, and 10% the left hand. We used the large, general population cohort of the UK biobank (~500,000 participants) to study possible relations between early life factors and adult hand preference. The probability of being left-handed was affected by the year and location of birth, likely due to cultural effects. In addition, handedness was affected by birthweight, being part of a multiple birth, season of birth, breastfeeding, and sex, with each effect remaining significant after accounting for all others. Maternal smoking showed no association with handedness. Analysis of genome-wide genotype data showed that left-handedness was very weakly heritable, but shared no genetic basis with birthweight. Although on average left-handers and right-handers differed for a number of early life factors, all together these factors had only a minimal predictive value for individual hand preference. Therefore other, unknown effects must be involved, including possible environmental factors, and/or random developmental variation with respect to the left-right formation of the embryonic brain.

**Significance statement:** Left-right laterality is an important aspect of human brain organization which is set up early in development. Left-handedness is an overt and relatively prevalent form of atypical brain laterality. Various, often related, early life factors have been previously studied in relation to handedness, but often in small samples, or samples with biased selection schemes. Here we have performed the largest ever study of left-handedness in relation to early life factors. Left-handedness was very weakly heritable and there were significant effects of various factors such as birthweight, which remained significant after controlling for all others. However, considered all together, early life factors still had poor predictive power for the handedness of any given individual. Very early developmental perturbations, caused by environmental or chance effects in embryonic development, are therefore likely to cause left-handedness.

## Introduction

Roughly 90% of people have a preference for using the right hand for complex manual tasks (1-3). A minority of roughly 10% prefer to use the left hand, and a smaller group of roughly 1% has no clear preference, the so-called ‘ambidextrous’ people. As a striking human behavioural polymorphism, handedness has attracted a lot of attention in both the scientific and popular literature. For example, personality traits and cognitive skills have been claimed to associate with handedness (4, 5). The prevalence of non-right-handedness has also been found to be increased in people with various cognitive or psychiatric disorders (6, 7).

Hand preference becomes established within the first two years of life, but prenatal observations using ultrasound scanning have indicated an earlier initiation of the trait (8, 9). Gene expression analysis has revealed left-right differences in the human central nervous system as early as four weeks post conception (10), which indicates that laterality is an innate and pervasive property of the brain. The strong skew towards right-handedness at the population level suggests that right-hand-preference is the typical or default arrangement for humans, while left-handedness may result from genetic, environmental or random perturbations that influence the central nervous system during early development (although alternatives to this view have been discussed (11, 12)).

One biological effect on handedness is known to be sex, with males more likely to be left-handed than females (2, 13). For example, in a U.S. dataset aged 10-86 years, the proportion of non-right-handers among 664,114 women was 9.9%, versus 12.6% among 513,393 men (2). Previous studies have also shown that genetic variation contributes modestly to left-handedness, with heritability estimates ranging from 0.03 for SNP-based heritability in the UK Biobank (N>500,000) (14), to 0.25 in twin studies (15, 16), and even > 0.5 in small family studies (17, 18). A number of candidate genes or genetic pathways have been proposed to be involved with varying degrees of statistical genetic support (19-22), but no genetic mechanisms or biological processes have yet been implicated unambiguously. In addition, no clear markers of brain anatomical asymmetry have been found to associate with handedness (23).

One of the problems with assessing handedness is that, historically, people who are not right-handed have often been made to use their right hand for writing, handling cutlery, and various occupational tasks (2, 24). As a consequence, a proportion of otherwise left-handed or ambidextrous people has become right-handed, while possibly also a number of left-handed people have become ambidextrous through this enforcing (25). The rate of enforced right-handedness varies between cultures (26), but has typically shown a decline over recent decades: in many countries, proportions of left-handers have increased with time, probably because society has become more tolerant of variation (2, 27, 28).

Among the early life factors that have been studied for associations with hand preference are the month of birth (29-31), being part of a multiple birth (32-35), birthweight (11, 15, 36), breastfeeding (37), and maternal smoking (38, 39). Effects of birthweight and multiple birth seem generally consistent throughout the literature; for example a recent study of two datasets of triplets, each numbering roughly 1000 participants, showed that lower birthweight was associated with non-right handedness (35). However, other effects remain equivocal. For example, previous studies have sometimes not taken the sex or age of participants into account, or have not accounted for the country of origin, so that the analyses may have been partly confounded. Other studies have only considered university students or other convenient or biased sampling selections, which may have resulted in an incomplete picture.

A very large, and well characterised, population-based cohort such as the UK Biobank, which includes hundreds of thousands of participants, allows multiple potential factors to be considered together, while providing unprecedented statistical power to begin to disentangle them. In this study, we analysed a number of early life factors that might influence adult hand preference in the UK Biobank dataset. In addition, as genome-wide association data are available for this cohort, we were able to assess the genetic correlations between handedness and these other factors, some of which are heritable in their own right. Genetic correlation is a measure of the extent to which the same genetic variation, over the entire genome, affects two traits.

## Results

### Factors associated with left-handedness

Data were obtained from the UK Biobank cohort, which is an adult population cohort (40). In total, the dataset comprised 501,730 individuals (Table 1), but exclusions for high residual genetic relatedness (see Methods) left 421,776 individuals for whom demographic information (year of birth and sex) is illustrated in Figure S1, and further drop-out then varied according to the availability of each specific variable (see Tables 3 and 4; for example, information on birthweight was available for 62% of the females and 47% of the males).

The distribution of answers to the hand preference question is shown in Table 1. The ambidextrous group was found to be inconsistent in their answers across timepoints (Methods), so that we focussed here only on the binary trait of left-handedness versus right-handedness.

**Table 1.** Distribution of responses to question about handedness

| Hand use | Males | Females | Total |
| --- | --- | --- | --- |
| Right-handed | 199,915 (87.4%) | 246,021 (90.1%) | 445,936 (89%) |
| Left-handed | 23,792 (10.4%) | 23,059 (8.4%) | 46,851 (9.3%) |
| Use both right and left hands equally | 4,847 (2.1%) | 3,813 (1.4%) | 8,660 (1.7%) |
| Prefer not to answer | 169 (0.007%) | 114 (0.004%) | 283 (0.005%) |
| TOTAL | 228,723 | 273,107 | 501,730 |

A number of early life variables were available in the UK biobank data. Table 2 shows the measures that were available. Month of birth was modelled using a cosine function to represent a continuous seasonal effect (Methods). As the degree of cultural enforcing of right-hand use is known to have varied by year and country (see Introduction), we also included country of birth and year of birth as possible predictor variables. Finally, we included sex.

**Table 2.** Variables included in the analysis. See table 3a for sample sizes.

| Description | header | Type | Note |
| --- | --- | --- | --- |
| Country of birth | f.1647.0.0 | categorical | 4 UK countries, Republic of Ireland, Elsewhere |
| Breastfed as a baby | f.1677.0.0 | categorical | 1=Yes,0=no, -1=do not know, -3=prefer not to answer |
| Part of a multiple birth | f.1777.0.0 | categorical | 1=Yes,0=no, -1=do not know, -3=prefer not to answer |
| Maternal smoking around birth | f.1787.0.0 | categorical | 1=Yes,0=no, -1=do not know, -3=prefer not to answer |
| Birthweight | f.20022.0.0 | continuous | (kg) |
| Sex | f.31.0.0 | categorical | 0=Female,1=Male |
| Year of birth | f.34.0.0 | continuous (integer) | between 1934 and 1971 |
| Month of birth | f.52.0.0 | continuous* | 12 months |

\*transformed before analysis, see main text

In univariable analyses, a higher probability of being left-handed was associated with being male, being part of a multiple birth, not being breastfed, having lower birthweight, being born in a more recent year, and being born in summer (all p-values < 0.05; Table 3 for categorical variables, Table 4 for continuous variables). The association of year of birth and left-handedness is shown in Figure S2, that of birthweight and left-handedness in Figure S3, and month of birth and left-handedness in Figure S4. The different countries within the UK also differed in rates of left-handers, with Wales having the lowest proportion, and people who were born outside the UK even lower (Table 3).

In separate univariable analyses of males and females (Table S1 and S2), the cosine function of month of birth only had an effect in females (Figure S4, p=0.388 in males, p=5.6e-05 in females).

**Table 3.**
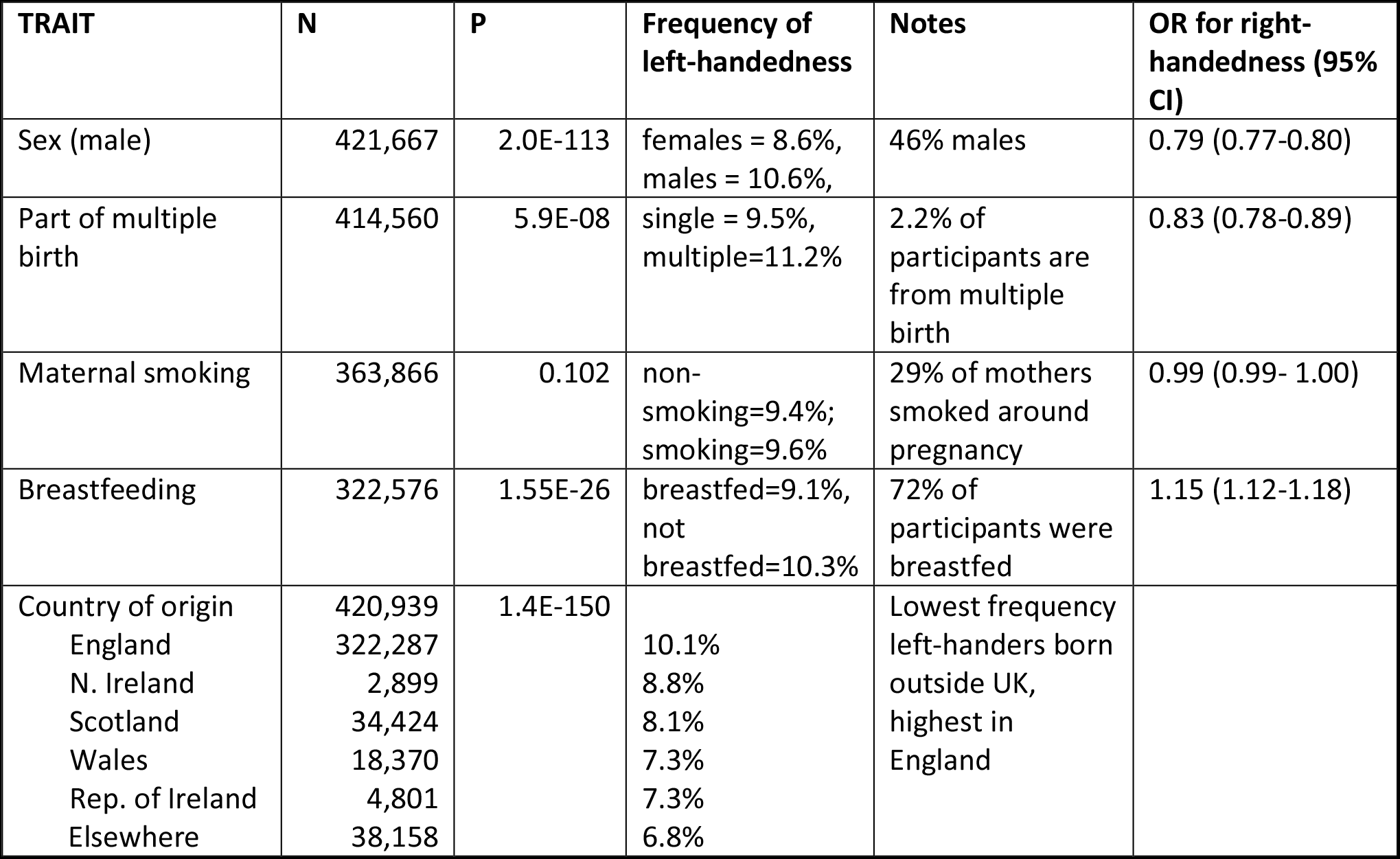
Univariable analysis of categorical early life variables and handedness. OR refers to the odds ratio, CI to the confidence interval.

**Table 4.** Univariable analysis of continuous early life variables and handedness

| TRAIT | N | p-value | Note | Effect on logit Right Handedness |
| --- | --- | --- | --- | --- |
| Year of birth | 421,667 | 1.0E-30 | Increase ~ 0.7percentage -points per decade | -0.007yr<sup>-1</sup> |
| Birthweight | 231,155 | 0.0009 | Left-handers are ~26g lighter on average | 0.035kg<sup>-1</sup> |
| Cosine(month) | 421,667 | 0.004 | See Figure S4 | 0.011 |

### Multivariable modelling and the relations between predictor variables

All predictor variables having shown nominally significant (p<0.05) effects on handedness in univariable testing (i.e. all but maternal smoking) were then included in multivariable analysis, using general linear modelling (Methods), with right-handed/left-handed as the dependent variable. In the multivariable model including both sexes, the probability of being left-handed was influenced by sex, year of birth, birthweight, country of birth, multiple birth, breastfeeding and the cosine function of month of birth (all p<0.05) (Table 5). As the model fitting involved simultaneous entry, the significance of each of these variables indicates an independent effect after accounting for all others. All variables together significantly explained variation in handedness (p=2E-127), but the predictive power for individual handedness was low (pseudo R^2^ MacFadden =0.005).

Although tests for variance inflation showed that there was no distorting collinearity in the model, with all inflation factors below 1.2, most predictor variables were correlated or associated with each other to a degree in pairwise univariable testing (see Figure 1). For example, those from multiple births reported being born considerably lighter than singletons (2.46 kg vs 3.36 kg, p<2.2e-16), while breastfed children were born heavier than non-breastfed children (3.39 kg vs 3.25 kg, p<2.2e-16). In fact, birthweight was associated with all of the other variables (Figure 1), apart from the year of birth (p=0.75, Figure S5). As regards birthweight and month of the year, the heaviest children were born in September-October (Figure S6). Children from smoking mothers were born a little lighter than from non-smoking mothers (3.28 kg vs 3.37 kg, p<2.2e-16). Males were heavier on average than females at birth (3.45 kg vs 3.25 kg, p<2.2e-16), but still males showed a higher probability of left-handedness than females in the multivariable model, i.e. opposite in direction to the association of birthweight and handedness. Also sex was associated with a number of the other variables (Figure 1). Year of birth was not correlated with birthweight, but was associated with most of the maternal behavioural traits (Figure 1).

**figure 1.**
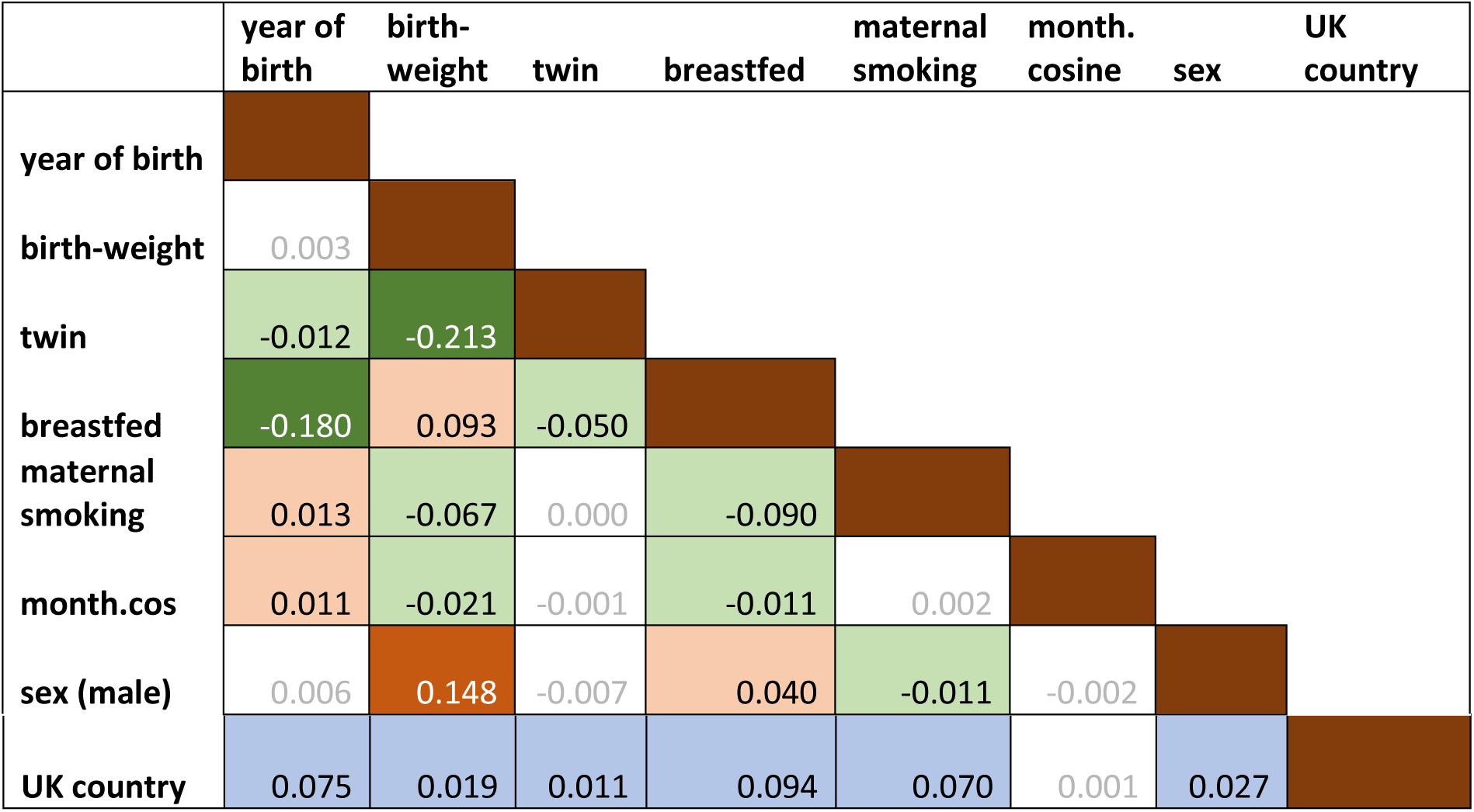
Associations between predictor variables. For associations between categorical variables, Cremer’s V is presented. Associations between continuous variables are shown as Pearson R. Associations between binary categorical and continuous variables are shown as Spearman rho. Associations between multi-category variable UK Country and continuous variables are shown as the ANOVA adjusted R. Colour and sign show the direction of the association between two binary variables, between two continuous variables or between binary and continuous variables (orange positive, green negative). Grey font indicates non-significant associations (p>0.001).

Despite the associations between the predictors, all predictors had independent effects on handedness in the multivariable model (Table 5). In the multivariable models that were fitted separately for males and females, again all included predictor variables were significant at p<0.05 (Tables S3 and S4), although month of birth and year-squared were not included in the model for males, as these were not significant in univariable testing in males only (P>0.05).

**Table 5.**
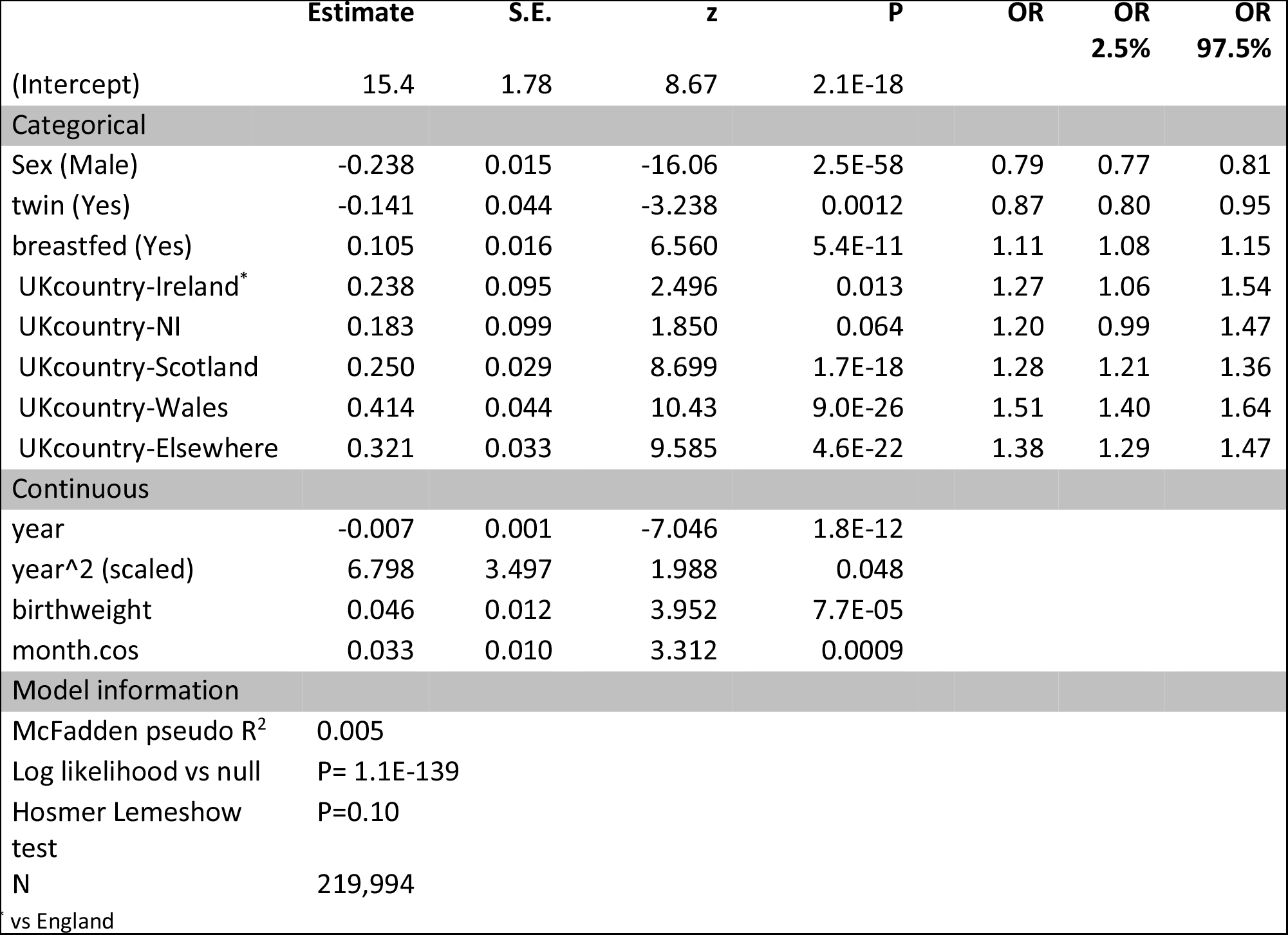
Multivariable logistic model for right-handedness, all participants

### Heritability and genetic correlation

SNP-based heritability is a measure ranging from 0 to 1 which indicates the extent to which variation in a trait is influenced by the combined effects of variation at SNPs distributed over the genome (41). Handedness, birthweight, and being breastfed were previously reported to have low but significant SNP-based heritabilities in the UK biobank dataset (handedness 1.8% (se=0.00737), birthweight: 12%(se=0.006); breastfed: 4.5% (se=0.00674) (42). Because we found that the latter two variables were associated with handedness in the present study (see above), we re-calculated their SNP-based heritabilities and then measured their genetic correlations with handedness, which had not been investigated before. Genetic correlation analysis measures the extent to which variability in a pair of traits is influenced by the same genetic variations over the genome.

Consistent with a previous analysis of the UKBiobank dataset (42), we found low but significant SNP-based heritabilities of left-handedness (4.35%), birthweight (15.47%), and being breastfed (5.94%) (see supplementary table S5). The analysis of genetic correlation between these measures was novel to the current study (Table S6), but there was no significant genetic correlation between handedness and being breastfed, nor between handedness and birthweight (Table S6; although note that we had limited power to detect a genetic correlation below 0.1 between handedness and the binary trait of being breastfed (Figure S7)).

## Discussion

### General observations

In this study we assessed various early life factors in relation to the probability of becoming left-handed. The large and well characterized dataset provided by the UK Biobank allowed the detection of very subtle associations, as well as the power to test for residual effects of the individual factors after correction for all others. We confirmed a number of previously reported early life factors that influence handedness, which we discuss in detail further below. Being male was associated with left-handedness, as has been widely reported and discussed before (see Introduction)(2, 13). In addition, we confirmed a very low heritability for left-handedness, but found no genetic correlation with birthweight or being breastfed.

However, perhaps the most striking finding from our study is that, even when taken all together, the studied factors had only a tiny predictive effect for individual handedness. The biological basis of left-handedness therefore remains largely unexplained. It remains possible that some major, early life influences on handedness do exist, but which were not assessed in the UK Biobank dataset, and do not correlate strongly with any of the early life variables that were available. However, other possibilities are also plausible, which are not necessarily mutually exclusive. Firstly, handedness might be influenced by an accumulation of many, very small environmental influences, possibly having an effect during prenatal stages. What such environmental effects may be is currently unknown. Secondly, heterogeneous and rare genetic mutations may also be involved (43, 44), whose effects are not well captured by measures of SNP-based heritability, as the latter approach is focussed primarily on more common genetic variation (41).

A random model of early embryonic development is also compatible with our observations. For example, if the brain’s left-right asymmetry arises from only a subtle left-right bias in the early embryo, such as a gene expression gradient that has a lateralized mean across embryos, but a variance that spans the point of symmetry, then a minority of embryos would experience a reversal of the foundational cues for left-right brain patterning. Subsequent steps in development might then reinforce upon the original cue, and result in the bimodal trait of hand preference in adults. Assuming that hand dominance and/or division of labour between the hands is beneficial for fine motor control, then the fact that we have two hands essentially imposes a binary choice on a developmental program which may be more continuous in its original nature. The fact that human brain embryonic gene expression has been shown to be only very subtly lateralized is consistent with such a model (45, 46).

### Environmental effects

Notwithstanding the subtle associations of predictor variables with handedness that we found, some of these associations are consistent with the previous literature and relevant to remark on. Year and country of birth were among the strongest effects. The proportion of left-handers increased almost linearly with year of birth up to 1970, i.e. the birth year of the youngest participants, with ~ 0.7 percentage-points per decade (Figure S2). We attribute this to a decline in enforcing right-handedness, as has been discussed before (27, 28, 47), rather than reduced survival of non-right-handers (48). However, as noted above, it is also possible that unknown environmental influences are involved in handedness, especially prenatally, which might have changed over the decades.

As regards country of birth, while the average proportion of left-handers among people born outside the UK was 6.8%, it was 10.1% in England, and intermediate in the other UK countries. These differences between countries are likely to reflect mainly cultural effects. For example, forced hand switching during childhood may have been more prevalent outside of England, or may have continued for longer.

An effect of the cosine-transformed month of birth on hand preference was found in women, such that left-handedness was associated with being born in the summer. The effect of season of birth on hand preference has been unclear in the literature. In a number of studies, a stronger seasonal effect was found in males than in females (49, 50). In other studies, more left-handers were found among children born in March-July (29, 30, 51), but in other studies in winter (49, 50, 52). In yet other studies, no effect of season was detected (31, 53-55). In the UK biobank, we observed that birthweight varied with season, with a pattern that was similar in males and females: the highest average birthweight was in September-October and lowest in February (Figure S6) (56). However, in the multivariable model we tested for a residual effect of the month of birth after correction for birthweight, and month of birth in females remained significant. Given the conflicting results across various studies, and the subtle and sex-limited effect reported here, this putative effect on hand preference remains tentative.

### Additional early life factors

We confirmed that having a higher birthweight, not being part of a multiple birth, and being breastfed, all increase the probability of being right-handed, consistent with previous literature (see Introduction). Birthweight is a complex trait which reflects not only healthy variation but also non-optimal development or pathology. Insofar as lower birthweight was associated with left-handedness, this suggests that a minority of left-handers may be linked etiologically to developmental insults, as has been discussed elsewhere (12, 57, 58). As regards multiple birth, in a previous study, the effect of twinning was no longer detectable after accounting for the effects of birthweight and APGAR (**A**ppearance, **P**ulse, **G**rimace, **A**ctivity, **R**espiration) score (34). We saw a significant effect of multiple birth on handedness after correction for birthweight, but the UK Biobank includes no APGAR scores, so we could not assess the relevance of this additional variable. Note also that the UK Biobank variable ‘Part of a multiple birth’ makes no distinction between twins, triplets, quadruplets etc., although the large majority are expected to be twins.

Interestingly, the postnatal behaviour of breastfeeding was associated with right-handedness and was also positively associated with birthweight: non-breastfed children were lighter at birth. The probability that mothers breastfeed their children may, among other things (59), be associated with mother or baby health, which in turn may be partly reflected in birthweight. Even after accounting for birthweight, a significant association of breastfeeding with right-handedness remained, as has been found before (Denny, 2011). Whether this is due to an underlying prenatal factor that affects both handedness and breastfeeding, or a post-natal behavioural effect, cannot be inferred from the UK Biobank data.

With regard to maternal smoking around the time of birth, we found that this was not significantly associated with left-handedness. An effect of maternal smoking on handedness had been reported suggestively before (38), but not found consistently in all studies (39).

In addition to the factors investigated in this study, other early life factors have been reported in the literature, including birth order (49, 60-62), prenatal testosterone exposure (63-65), maternal age (49, 53, 66, 67), maternal stress during pregnancy (11, 68), and birth events such as caesarean delivery or prolonged labour (12, 57, 69). Though not all studies have found significant effects of these variables, a general interpretation of the literature is that less benign conditions are associated with higher proportions of left-handedness. Some of these factors may partly influence handedness through effects via birthweight. For example, second and third births were reported to result more often in right-handed children than first births, and births subsequent to third (49, 60, 67), while more left-handers were reportedly born to relatively young mothers or older mothers, than to mothers of intermediate age (67, 70). Birthweight first increases with maternal age and subsequently decreases (71, 72), and low-weight children and preterm births are more common among young (< 20) and older (> 30) mothers than mothers aged in-between (71, 73). Birth order necessarily correlates with maternal age, with births two and three occurring more often in the intermediate age range. However, birthweight has been shown to vary with birth order even after correction for maternal age (72, 74). Unfortunately, information on maternal age and birth order were not available for the UK Biobank dataset.

### Heritability and genetic correlation

We observed a weak SNP-based heritability for left-handedness (4.35%) which was consistent with previous reports, but there was no genetic overlap between handedness and birthweight or being breastfed. For handedness and birthweight, we had 80% statistical power to detect a genetic correlation as low as 0.18, so that the phenotypic correlation between handedness and birthweight that we observed is likely due to an underlying environmental cause, rather than genetic factors. However, it is well established that SNP-based heritability can only capture a proportion of total heritability, i.e. which is caused by common polymorphisms tagged on genotyping arrays (41). In large twin studies, the heritability of handedness was higher, around 20-25% (15, 16, 25). The same was the case for birthweight, which had a twin heritability ~25% or higher (75, 76). Therefore genetic effects mediated by rare genetic variation, which was not well captured in this dataset, may also be relevant to the heritability of handedness and birthweight, and in some cases might link these two traits.

### Limitations

The UK Biobank participants are older than the general population (birth years between 1934 and 1971), so that some effects in this cohort may be quantitatively or qualitatively different in younger cohorts (77). The UK Biobank cohort, ranging over 30 years in age differences, and collected cross-population, is also more heterogeneous than some other previously investigated cohorts for handedness. This may make our results broadly applicable to the general population, but on the other hand, it may mean that some effects were obscured by variation in factors that were not accounted for.

All variables, except sex and year of birth, were self-reported. This may introduce inaccuracies as recall may be imperfect. Also, it is possible that cultural differences affect recall, for example between geographical regions, different ages, or between the sexes. This may have reduced the estimates of effects of early life factors on handedness, compared to if they had been recorded from direct observation.

The present study treated handedness as a categorical trait, which is supported by the bimodal distribution of overall hand preference when compiled across a number of tasks, and its robust test-retest repeatability (78-80). However, some aspects of handedness might be more accurately defined by degree and not category.

We allowed for possible non-linear effects of continuous predictor variables in our analyses, but did not include interaction terms between predictors in our multivariable models, in order to avoid collinearity, overfitting, and very extensive multiple testing. As regards sex, we performed some analyses separately within the two sexes to allow for potentially different effects, and we presented some descriptive comparisons between the sexes, but again did not test formally for interaction effects that involve sex. The male and female cohorts differed in a number of respects. For example, the proportion of males was not constant across the years of birth (Figure S1), while women more often than men originated from the Republic of Ireland or elsewhere outside of the UK (Tables S1 and S2), and a much larger proportion of women than men reported their birthweight. Future hypothesis-driven work may investigate specific potential interactions of the various factors studied here.

## Methods

Data were obtained from the UK Biobank cohort, as part of research application 16066, with Clyde Francks as the principal applicant. The data collection in the UK Biobank, including the consent procedure, has been described elsewhere (40). Informed consent was obtained by the UK Biobank for all participants. For this study of early life factors we used measurements taken during the first visit, i.e. variables identified in the database by 0.0. In total, data were available for 501,730 individuals (Table 1). To avoid using non-independent data, we excluded randomly one individual from each pair of participants whose genetic relatedness was inferred to be 3rd degree or closer, on the basis of genotype data at single nucleotide polymorphisms (SNPs) spanning the genome, as previously calculated by Bycroft et al. (81). This left 421,776 individuals.

For a subset of 9,856 participants, answers to the question about hand preference were available for both the initial (0.0) and the second follow-up visit (2.0). We compared these answers for consistency. While right-handed and left-handed persons were mostly consistent (only 0.7% and 2.6% changed their answer respectively), out of 156 people who had initially answered “use both right and left hands equally”, 64 (41%) gave a different answer during follow-up. We therefore excluded all people who answered “use both right and left hands equally” at their first visit from further analyses, as well as the small number of people who had ticked ‘Prefer not to answer” (Table 1).

The early life variables which were available for this study are shown in Table 2, as well as sex, year and country of birth, which were also used as predictor variables for left-handedness. All variables were self-reported, except sex and date of birth (see UK Biobank Showcase; http://www.ukbiobank.ac.uk/), although sex information could be updated by the participants.

The entries “do not know” and “prefer not to answer” for all variables were treated as missing values.

As months of the year are not independent categories (neighbouring months are more similar to each other with respect to e.g. temperature and day length), one approach is to model the effects of season as a waveform function of the month (82). We followed this approach and tested:

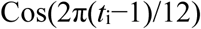

where *t_i_* is an integer from 1 to 12 representing the month of birth. This cosine function has extremes in summer and winter.

We excluded individuals with birthweight heavier than 6.0 kg to avoid outlier effects.

### Statistical analysis

All statistical analyses were performed with Rstudio, using R version 3.4.0.

#### Univariable analysis of categorical predictors of handedness

Associations between handedness and each of the categorical variables (country of birth, breastfed, multiple birth, maternal smoking, sex) were investigated with chi-square tests of independence.

#### Univariable analysis of continuous predictors of handedness

For testing univariable associations between handedness and continuous variables (birthweight, year of birth and cosine of month), logistic regression was used. In addition, univariable effects on proportions of left-handed people were visualised to assess whether non-linear relations were playing a role (Figures S2, S3). A model including either birthweight squared, or year of birth squared (as orthogonal vectors created by R function poly() from the ‘stats’ package), was compared to the corresponding model with the single variable to establish whether the squared predictor made a significant additional contribution.

The above analyses of handedness were also carried out separately within the two sexes.

#### Multivariable analysis of handedness predictors

For multivariable analysis, glm (general linear model) was used in R v3.4.0, for the binomial family of models. Participants with missing values for any of the predictor variables were excluded. The threshold for significance in the multivariable model was set at 0.05, i.e. testing whether each variable made a contribution beyond the combined effects of all others, in simultaneous entry. Collinearity was checked with the VIF (Variance Inflation Factor) function in R. Model fit was estimated with the Hosmer-Lemeshow test, using 15 quantiles, while the log likelihood of the full model vs the null model (with no predictors for handedness) was also estimated. In addition, the McFadden pseudo R2 was computed.

In the multivariable model, 219,994 participants without missing values were included: 83,506 males and 136,488 females. Multivariable analysis was also repeated for males and females separately.

#### Further statistical analysis

We investigated the pairwise relations between predictor variables as follows: For categorical pairs of variables the Chi square test was used to calculate Cramer’s V (i.e. a statistic scaled from 0 to 1 as an indication of the degree of non-independence). The R command *assocstats* was used for these calculations. For continuous pairs of variables the Pearson correlation coefficient R was calculated. When one of a pair of variables was dichotomous and the other continuous, Spearman’s rho was calculated. For Country of Birth in relation to continuous variables, ANOVA was used in which the adjusted R was calculated.

### Genetic analysis

For this purpose, in addition to removing one from each pair of related subjects (see above), participants were also excluded when there was a mismatch of their reported and genetically inferred sex (N=378), putative aneuploidy (N=652), excessively high genomewide heterozygosity (> 0.19) or genotype missingness (missing rate >0.05) (N=986), and we also restricted the analysis to participants with British ancestry as used by Bycroft et al (81). After this genetic quality control, there were a maximum of 335,998 participants per variable (see table S5 for sample sizes per variable).

We then calculated the SNP-based heritabilities and genetic correlations between two traits using Restricted Maximum Likelihood estimation implemented as –reml in BOLT-LMM (v2.3)(83). We used a genetic relationship matrix that included 547,108 genotyped SNPs (Minor Allele Frequency (MAF) >1% and genotyping rate across subjects >99%), and the pre-computed linkage disequilibrium (LD) scores based on 1000 Genomes European-descent data (https://data.broadinstitute.org/alkesgroup/LDSCORE/). The top ten principal components capturing genetic diversity in the genome-wide genotype data, calculated using fastPCA (84) and provided by the UKBiobank (81), were included as covariates to control for population structure, as well as sex, age, genotyping array, and assessment centre. For the binary traits handedness and being breastfed, the population and sample prevalence were assumed to be the same, in order to transform the SNP-based heritabilities to the liability scale using the R code provided by Pulit et al (85).

Power analyses for assessing genetic correlations between handedness and the two other traits (i.e. birthweight and being breastfed) were performed using the GCTA-GREML power calculator (86), considering the relevant sample sizes, SNP-based heritabilities and trait prevalences (Table S5). Since BOLT-REML heritability estimates and standard errors are close to GCTA-REML estimates (87) this calculator gives an indicative estimate. Results of the power analysis are shown in Figure S7.

## Manuscript information

### Ethics statement

This study utilized deidentified data from the baseline assessment of the UK Biobank, a prospective cohort study of 500,000 individuals (age 40–69 years) recruited across Great Britain during 2006– 2010 (40). The protocol and consent were approved by the UK Biobank’s Research Ethics Committee.

## Author contributions

CGFdK: Conceptualization, Formal Analysis, Methodology, Visualization, Writing – Original Draft Preparation. ACC: Data Curation, Formal Analysis, Writing – Original Draft Preparation. CF: Conceptualization, Funding Acquisition, Project Administration, Supervision, Writing – Review & Editing

## Supplemental

Supp. Figure 1 Male and female participants by year of birth

Supp. Figure 2 Left-handedness vs year of birth

Supp. Figure 3 Left-handedness vs birthweight in males and females

Supp. Figure 4 Left-handedness vs month of birth

Supp. Figure 5 Birthweight vs year of birth

Supp. Figure 6 Birthweight vs month of birth

Supp. Figure 7. Power analysis for genetic correlation

Supp. Table 1. Univariable analysis of early life variables and handedness in males

Supp. Table 2. Univariable analysis of early life variables and handedness in females

Supp. Table 3 Multivariable model for right-handedness, male participants

Supp. Table 4 Multivariable model for right-handedness, female participants

Supp. Table 5. Heritability estimates of handedness and covariates of interest

Supp. Table 6. Genetic correlation of handedness with a second trait

